# Climatic suitability and invasion risk of the elm zigzag sawfly in North America

**DOI:** 10.1101/2025.08.21.671666

**Authors:** Claudia Nuñez-Penichet, Zenia P. Ruiz-Utrilla, Daniel Rojas-Ariza, Weverton C. F. Trindade, Joanna L. Corimanya, Andres Herrera, Kier Mitchel E. Pitogo, A. Townsend Peterson

## Abstract

The elm zigzag sawfly, *Aproceros leucopoda* (Hymenoptera), native to eastern Asia, is among the most concerning potential pests for elm trees (*Ulmus* spp.); elms form an important component in many North Temperate forests. This sawfly species invaded Europe in 2003 and spread rapidly across much of that continent. In 2020, it was recorded in North America, and it has since become established in several parts of the eastern United States and Canada. Sawfly infestations can cause severe elm tree defoliation, branch die-back that weakens tree health, and potentially tree mortality. Sawfly invasions have the potential to further exacerbate elm decline, especially in conjunction with other pressures, including Dutch elm disease (*Ophiostoma ulmi*). We used rigorous approaches from distributional ecology to explore climatic suitability for *A. leocopoda* across North America, considering various sources of uncertainty in the data. We found that, without control, the elm zigzag sawfly could establish populations across eastern Canada, much of the central-eastern and northeastern United States, as well as in the Pacific Northwest. More southern areas of North America were not climatically suitable for this species. Predicted suitable areas for the sawfly overlap broadly with elm distributions, highlighting the need to control this invasion to mitigate potential economic and environmental impacts.

## Introduction

Biological invasions rank among the most significant drivers of biodiversity loss, causing changes in community structure and composition [1–4]. The negative impacts of invasive alien species (IAS) extend to ecosystems, biodiversity, human health, and the economy [3,5,6]. One of the most recent introductions in North America is the hymenopteran *Aproceros leucopoda* (elm zigzag sawfly; Hymenoptera: Argidae), Takeuchi, 1939. *Aproceros leucopoda* is a multivoltine species native to East Asia, capable of producing 4-6 generations per year [7,8]. Short-lived adults reproduce primarily through parthenogenesis, which is common in sawflies [9–11]. The entire life cycle, from egg laying to adult emergence, spans approximately 24-29 days [7]. Larvae of this species feed only on elm (*Ulmus* spp.) leaves, leaving a distinctive zigzag pattern of damage [7], which has caused severe defoliation in Europe [12,13]. The appearance of *A. leucopoda* in Europe began in Hungary and Poland in 2003 [7]; today, it is considered an exotic species in most European countries, particularly in Eastern Europe [14]. The defoliation rate of infested elm trees in Europe was 74-93% of leaves consumed by *A. leucopoda* larvae; in some cases, complete defoliation was reported [7,15]. Although elm foliage recovers post-damage [7], mortality risk increases with additional stressors like Dutch elm disease, elm yellows, and other diseases that target *Ulmus* [13].

This species was first reported in the Americas in 2020 in Québec, Canada, via the community science website, iNaturalist [16]. The succeeding years saw reports of the sawfly or signs of its presence (i.e., elm leaves with the characteristic zigzag damage pattern) in Virginia in 2021 and in Pennsylvania, North Carolina, Maryland, and New York in 2022 [8,17]. This species has since been reported hundreds of times in North America on iNaturalist, with records from most of southeastern Canada and the northeastern United States. The most recent reports of this species on the platform were in Wisconsin and Minnesota in August and September 2024, pointing to a rapid range expansion of this species across North America.

Elm genus *Ulmus* comprises 45 woody species distributed mainly in temperate and subtropical regions [18]. In the U.S., 39 elm species are reported [19], among which *U. americana* is classified as an endangered species by the IUCN [20]. These emblematic trees hold significant economic value in the U.S., given their widespread use as a construction material and their use as ornamental trees in urban landscapes [21,22]. Additionally, elms provide critical ecological services in North American floodplains [23]. In this context, establishment and further spread of *A. leucopoda* in North America, combined with other elm stressors such as Dutch elm disease, could severely impact elm survival, with rapid economic and ecological consequences [24]. Therefore, it is crucial to understand the potential distributional patterns of this invader in North America to be able to implement more efficient and timely preventive, control, and eradication measures.

The potential and actual distributional areas of a species in a geographic area are determined by the joint presence of suitable biological interactions (area **B**), favorable environmental conditions (area **A**), and the species’ capacity to access those suitable areas (area **M**; [25]). Characterizing these three factors for any given species can represent a challenge because, although several methods by which to estimate **A** are available, information on **B** and **M** is often much more limited [26]. This lack of information may lead to overprediction of a species’ potential distribution areas, as regions with suitable environmental conditions may still be inaccessible owing to geographic barriers and/or negative biotic interactions [27].

In the case of invasive alien species (IAS), human intervention often enables them to reach previously inaccessible areas [28,29], thereby augmenting the **M** area. When environmental conditions in the invaded area are favorable, successful establishment of the IAS depends on their biology and interactions with other species, facilitating rapid population growth [30,31]. Ecological niche modeling (ENM) is a valuable tool to assess and predict the potential areas susceptible to invasion by IAS, relying solely on climatic variables without considering the biological interactions or mobility factors [32–34]. This approach has been used to model recent invasions in North America, such as the cases of *Vespa mandarinia* and *V. velutina*, where potential invasion areas were predicted based on climatic data [35,36]. Identifying these potential invasion areas is key to developing management strategies, especially for short-lived, highly dispersive species like *A. leucopoda* [34]. Given the need for early detection and rapid response to prevent the spread and establishment of IAS, the aim of this study is to identify the potential areas of invasion by *Aproceros leucopoda* in North America and to assess its potential impact on *Ulmus* in this region.

## Methods

### Occurrence and environmental data

We gathered occurrence data for the elm zigzag sawfly in both its native and invaded ranges, as well as for the genus *Ulmus* in North America. The compiled dataset of *A. leucopoda* (2930 occurrences in total) included records from the Global Biodiversity Information Facility (GBIF; 2688 records, Dr. H. Hara personal communication (12 records), and literature sources (230 records). Records were most numerous from Europe (2218 records), and East Asian records were more scarce (41 records); only 727 records were available from North America. For *Ulmus*, we obtained 604619 occurrences from GBIF [19] and 224786 records from the Botanical Information and Ecology Network (BIEN; https://bien.nceas.ucsb.edu/bien/), totaling 829405 occurrences.

To download the occurrences from the online repositories, we used R version 4.3. [37], specifically the *rgbif* v. 3.7.9 [38] and *BIEN* v.1.2.6 [39] packages to retrieve data from GBIF and BIEN, respectively. We cleaned the data for both taxa by removing records without latitude and longitude information, those with coordinates of (0°,0°), duplicate records, and records with low coordinate precision (<2 decimals on their longitude or latitude; [40]). We then excluded records within a 5 km radius of one another to reduce spatial autocorrelation and prevent model overfitting [41]. In the elm dataset, we observed a strong bias in data density among countries, likely due to differences in sampling effort or data reporting. To account for these potential biases, we reduced the number of occurrences in countries with the highest sampling densities by standardizing record density across countries to match the density of the country with the fewest occurrences per km^2^ (i.e., El Salvador; e.g., [35]). After all cleaning steps, we retained a total of 707 occurrences for *A. leucopoda* and 2221 for *Ulmus* (Fig. 1).

**Fig 1.**
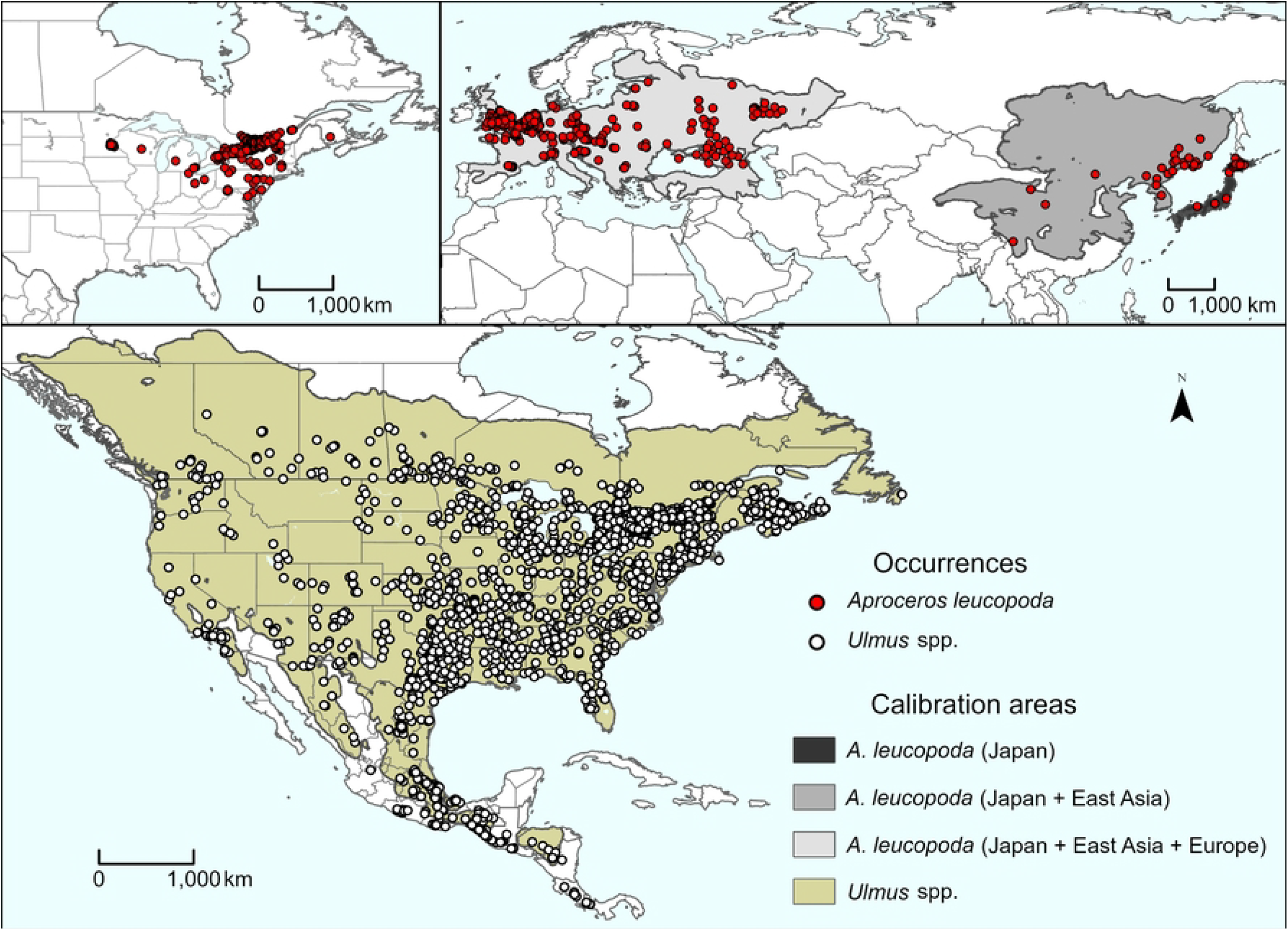
Occurrences and calibration areas (M) of the taxa used in this study. Top: *Aproceros leucopoda* in its native and invaded areas. Bottom: *Ulmus* spp. in North America

We obtained 19 environmental variables representing monthly temperatures and precipitations from WorldClim (v2.1, [42]; https://www.worldclim.org/data/worldclim21.html) at 2.5′ (∼5 km) resolution for the period 1970–2000. We excluded four variables (mean temperature of wettest quarter, mean temperature of driest quarter, precipitation of warmest quarter, precipitation of coldest quarter) owing to spatial inconsistencies [43]. For the sawfly, environmental variables were cropped to the Northern Hemisphere, *Ulmus* was considered in North America. To reduce collinearity between variables and reduce dimensionality [44], we performed a principal components analysis (PCA) on the cropped environmental variables, and retained the first five components (PCs), which accounted for 97% of the total variance (Table S1).

### Ecological niche modeling

We used clean datasets and the five selected PCs to create ecological niche models for both taxa, using Maxent [45]. We used 70% of the occurrence data for model training and the rest to evaluate model performance. We also created three calibration areas for the zigzag sawfly models, and one for calibrating elm models; these areas were defined based on dispersal simulations using the R package *grinnell* (v.0.0.22, [46]). These simulations were conducted under constant climatic conditions (see “stable framework” in [46] using a single set of environmental layers for the period of 1970–2000 (Worldclim v2.1). For the sawfly, we used the three areas depicted in Fig. 1, which were created using the occurrence data of the zigzag sawfly for each region and parameterized with the following settings: a starting proportion of 0.9 (i.e., 90% of the data used as starting points of the simulation), a normal dispersal kernel with 8 kernel spread values (i.e., 1-8), 5 numbers of dispersal events (i.e., 30, 60, 120, 180, 240), a maximum number 5 of dispersers per occupied cell, a suitability threshold of 5, and 10 simulation replicates. For *Ulmus*, we defined North America as the calibration area and applied a starting proportion of 0.9, a normal dispersal kernel with spread values of 1 to 8 pixels, 4 numbers of dispersal events (i.e., 20, 40, 80, 160), a maximum of 2 dispersers per occupied cell, a suitability threshold of 5, and 10 simulation replicates. We selected best simulation results, considering establishment of full contiguity across the area of simulation and stability of simulation results across different parameter settings, areas obtained with a value of 6 and 5 of kernel spread and 120 and 180 dispersal events for the sawfly and *Ulmus*, respectively.

We created a total of 1170 candidate models for *Ulmus* and 3510 for each calibration scenario used for the zigzag sawfly (see Fig. 1). All models were produced considering all combinations of the following parameter settings: 5 PCs, 5 feature classes (q, lq, lp, qp, lqp; l indicates linear, q quadratic, and p product), and 9 regularization multiplier values (i.e., 0.1, 0.25, 0.5, 0.75, 1, 2, 3, 4, 5). We evaluated model performance using partial ROC [47], predictive ability (omission rates at an acceptable omission rate *E* = 5%; [48]), and the Akaike Information Criterion corrected for small samples (AICc, for evaluating model fit vs model complexity, [49]). Model selection was done based on the statistical significance of partial ROC values, omission rates <5%, and ΔAICc values <2. Model calibration and selection were done using the *kuenm* R package [50].

Final models selected using the criteria described above (Table 1) were visualized across the calibration region and across all of North America. We performed 10 bootstrap replicates and explored three extrapolation types for each selected model: no extrapolation, extrapolation with clamping, and free extrapolation. When multiple models met the selection criteria, we applied a model consensus approach, using the median of single-model medians [35] to obtain a single final model for each calibration scheme. Finally, we converted models for both taxa into binary presence/absence maps using a 95% training presence threshold (*E* = 5%) and calculated their spatial overlap. We also visually evaluated the predictive performance of the *A. leucopoda* models by comparing the resulting predictions under each calibration scheme with the species’ known invaded areas in North America.

**Table 1.** Parameters selected for the final models for each of the three calibration areas for *A. leucopoda* and the *Ulmus* genus calibration area.

| Calibration area | Feature classes | Regularization multiplier | Variable set | Mean AUC ratio | P-value pROC | Omission rate (5%) | $\Delta AICc$ | Number of parameters |
| --- | --- | --- | --- | --- | --- | --- | --- | --- |
| *Ulmus* spp. |  |  |  |  |  |  |  |  |
| North America | 0.1 | l, q, p | 26 | 1.34 | 0 | 0.040 | 0 | 19 |
| *Aproceros leucopoda* |  |  |  |  |  |  |  |  |
| Japan | 0.10 | q | 4 | 1.73 | 0 | 0 | 0 | 2 |
|  | 0.25 | q | 4 | 1.72 | 0 | 0 | 0.03 | 2 |
|  | 0.25 | q, p | 4 | 1.74 | 0 | 0 | 0.10 | 2 |
|  | 0.50 | q | 4 | 1.73 | 0 | 0 | 0.13 | 2 |
|  | 0.75 | l, q | 4 | 1.73 | 0 | 0 | 0.30 | 2 |
|  | 0.75 | q | 4 | 1.72 | 0 | 0 | 0.28 | 2 |
|  | 0.50 | q, p | 4 | 1.73 | 0 | 0 | 0.39 | 2 |
|  | 0.50 | l, q, p | 4 | 1.73 | 0 | 0 | 0.39 | 2 |
|  | 0.10 | q | 7 | 1.69 | 0 | 0 | 0.46 | 2 |
|  | 1.0 | q | 4 | 1.73 | 0 | 0 | 0.47 | 2 |
|  | 1.0 | l, q | 4 | 1.72 | 0 | 0 | 0.47 | 2 |
|  | 0.25 | q | 7 | 1.70 | 0 | 0 | 0.50 | 2 |
|  | 0.50 | q | 7 | 1.70 | 0 | 0 | 0.62 | 2 |
|  | 0.75 | q | 18 | 1.70 | 0 | 0 | 0.80 | 2 |
|  | 0.75 | q | 7 | 1.70 | 0 | 0 | 0.80 | 2 |
|  | 0.75 | l, q, p | 4 | 1.72 | 0 | 0 | 0.80 | 2 |
|  | 0.75 | q, p | 4 | 1.74 | 0 | 0 | 0.80 | 2 |
|  | 1.0 | q | 18 | 1.70 | 0 | 0 | 1.03 | 2 |
|  | 1.0 | q | 7 | 1.70 | 0 | 0 | 1.03 | 2 |
|  | 1.0 | l, q, p | 16 | 1.73 | 0 | 0 | 1.30 | 2 |
|  | 1.0 | q, p | 16 | 1.74 | 0 | 0 | 1.30 | 2 |
|  | 1.0 | l, q, p | 4 | 1.73 | 0 | 0 | 1.30 | 2 |
|  | 1.0 | q, p | 4 | 1.74 | 0 | 0 | 1.30 | 2 |
|  | 2.0 | q | 16 | 1.74 | 0 | 0 | 1.55 | 2 |
|  | 2.0 | q | 24 | 1.74 | 0 | 0 | 1.55 | 2 |
|  | 2.0 | l, q | 16 | 1.74 | 0 | 0 | 1.55 | 2 |
|  | 2.0 | l, q | 24 | 1.74 | 0 | 0 | 1.55 | 2 |
|  | 2.0 | q | 15 | 1.74 | 0 | 0 | 1.55 | 2 |
|  | 2.0 | q | 4 | 1.74 | 0 | 0 | 1.55 | 2 |
|  | 2.0 | l, q | 15 | 1.73 | 0 | 0 | 1.55 | 2 |
|  | 2.0 | l, q | 4 | 1.73 | 0 | 0 | 1.55 | 2 |
| Japan +<br>East Asia | 0.10 | l, q | 13 | 1.76 | 0 | 0 | 0 | 5 |
|  | 0.25 | l, q | 13 | 1.76 | 0 | 0 | 0.05 | 5 |
|  | 0.50 | l, q | 13 | 1.70 | 0 | 0 | 0.22 | 5 |
|  | 0.10 | l, q, p | 13 | 1.74 | 0 | 0 | 0.57 | 9 |
|  | 0.50 | l, q | 23 | 1.74 | 0 | 0 | 0.80 | 6 |
| Japan +<br>East Asia<br>+ Europe | 0.50 | l, q, p | 21 | 1.53 | 0 | 0.045 | 0 | 12 |

### Model uncertainty

We assessed model uncertainty for both taxa related to the risk of strict extrapolation when projecting to non-analogous environmental conditions. To this end, we used the mobility-oriented parity (MOP) metric [51], which measures the similarity between the closest 5% of environmental conditions at background points in the calibration area and each environmental condition in the projection area. We calculated MOPs for each group of occurrences and calibration areas (North America (*Ulmus*), Japan; Japan + East Asia; and Japan + East Asia + Europe), using only the variables selected in the respective final models. Projection areas with zero similarity values compared to the calibration areas indicate non-analogous conditions and higher uncertainty. In these regions, predicted suitability results from strict extrapolation, and model predictions should be interpreted with caution. All MOP analyses were conducted in R using the *mop* package [52].

All data need it to replicate our results can be found on the https://figshare.com/articles/dataset/Data_for_the_manuscript_Climatic_suitability_and_invasion_risk_of_the_elm_zigzag_sawfly_in_North_America_/29824700 online repository. All code used in this study can be found at https://github.com/claununez/Sawfly-EMM.

## Results

### Ecological Niche Model Predictions

From the 1170 models created and evaluated for *Ulmus*, only one met all three evaluation criteria (omission rate, ΔAICc, and partial ROC). For *A. leucopoda*, by contrast, 31, 5, and 1 models out of the 3510 evaluated met the criteria for the calibration areas Japan, Japan + East Asia, and Japan + East Asia + Europe, respectively (Table 1).

The final model for *Ulmus*, using free extrapolation, identified likely suitable areas across most of Mexico and the United States. Unsuitable areas were primarily located in the central western region of the United States and the northern Mexican states of Sonora and Chihuahua (Fig. 2).

**Fig 2.**
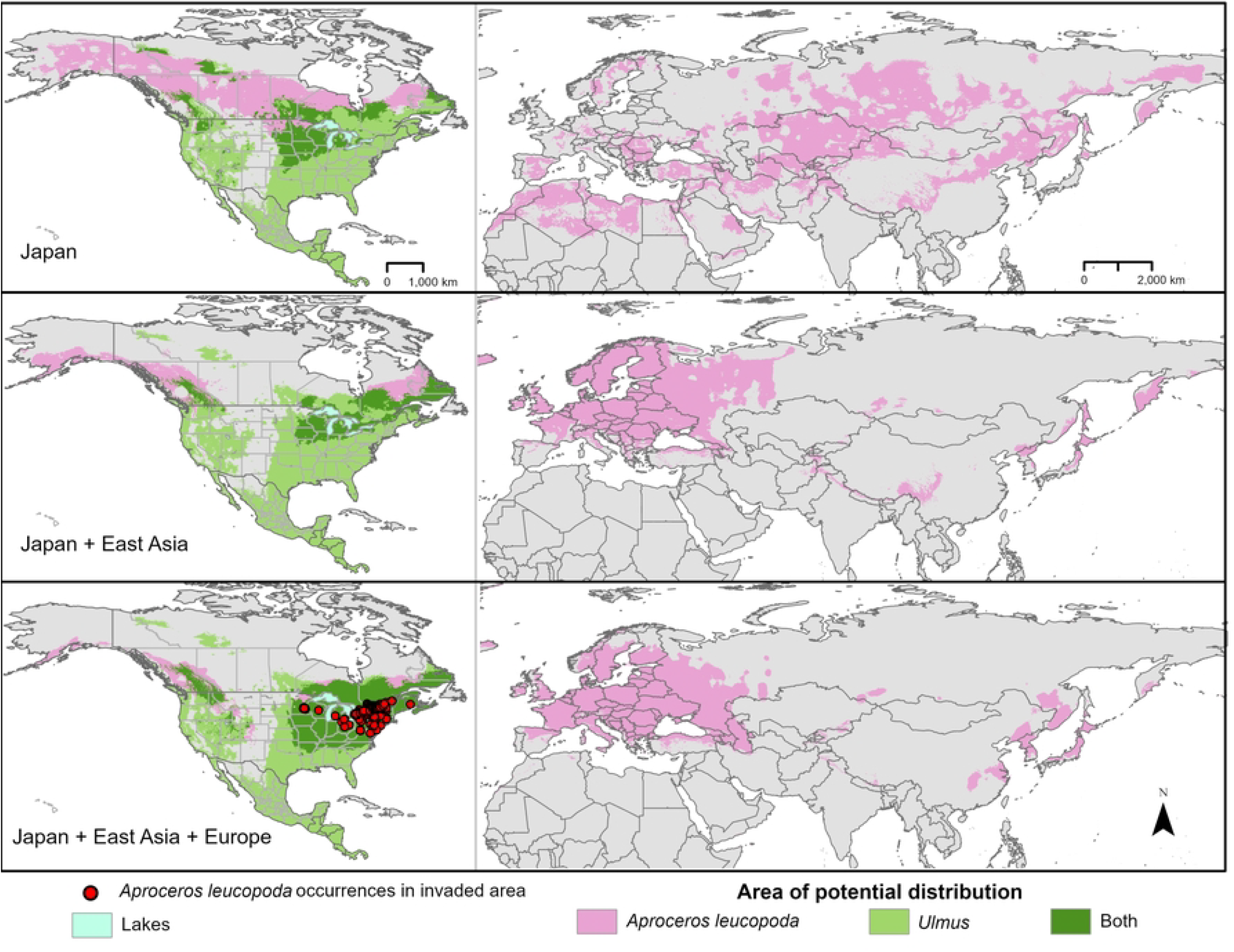
Areas of potential suitability of *Aproceros leucopoda* for models with free extrapolation

Predicted suitable areas for the zigzag sawfly (*A. leucopoda*) varied substantially depending on the calibration scheme. Models calibrated using only Japan failed to anticipate the species’ invasive potential in Europe. In contrast, models calibrated with Japan + East Asia successfully captured the invaded ranges in both Europe and North America (Fig. 2), and models calibrated based on Japan + East Asia + Europe successfully anticipated the invaded range in North America. The invasion pattern in North America was best characterized by the models calibrated with the most inclusive dataset—Japan + East Asia + Europe (Fig. 2).

In North America, the predicted suitable areas, based on calibration regions from Japan, East Asia, and Europe that overlap with the *Ulmus* model, include major parts of the eastern United States and Canada. Specifically, the model identifies suitable regions in southern Quebec and Ontario in Canada, as well as in the northeastern and southern United States. The southern distribution limit extends across northern South Carolina, Georgia, Alabama, Mississippi, Louisiana, and Texas. In the west, suitable areas are also predicted in the U.S. states of Idaho, Montana, and Washington, as well as in the Canadian provinces of British Columbia and Alberta (Fig. 2). The distribution of areas predicted to be suitable for both *Ulmus* and sawfly, as well as the areas of overlap, varied among the extrapolation types considered (Table S2, Figure S1, and Figure S2).

Areas identified as having a risk of strict extrapolation for one variable include only the most northern parts of Canada and southwestern Mexico in the model calibrated using Japan + East Asia + Europe. In contrast, when models were calibrated using only Japan, large areas of non-analog conditions across most of the projection regions were detected for multiple variables (Fig. 3). The Japan + East Asia models were intermediate in the degree and distribution of areas presenting non-analog conditions.

**Fig 3.**
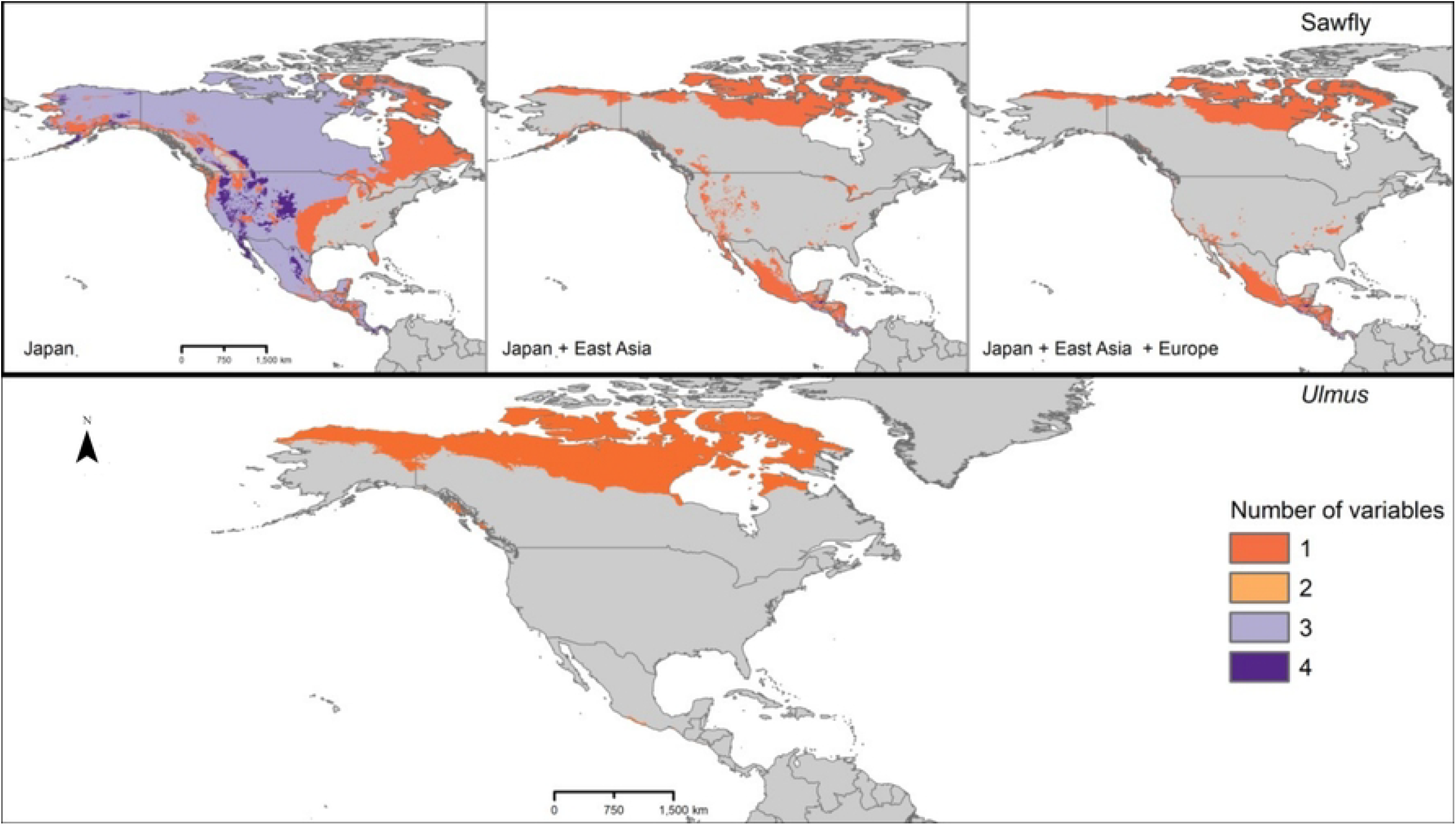
Areas identified as having a risk of strict extrapolation for the models of *Aproceros leucopoda* (top panels) and *Ulmus* spp. (bottom panel).

## Discussion

Managing invasions of alien species becomes increasingly challenging and resource-intensive in the later stages of the invasion process, such as once populations are established in the new areas [53]. Given that invasive alien species (IAS) drive biodiversity loss and cause significant ecological and economic impacts [54–57], predicting their invasion potential is crucial for development of biosecurity strategies [58,59]. Such predictions provide essential insights for developing cost-effective methods to mitigate early invasion stages [60] and for creating a “warning list” to enhance preventive measures in regions at risk [61,62].

Using ecological niche modeling (ENM), a method proven effective for predicting biological invasions [34,63–65], we estimated the potential distribution of *A. leucopoda* in North America. This work represents the first attempt to identify potential areas of invasion for *A. leucopoda* since its initial detection on the continent in the summer of 2020 [8]. Evaluating multiple calibration schemes allows us to assess the accuracy and limitations of models by providing a comparative framework for evaluating predictive performance.

Models calibrated using only the sawfly’s native range (Japan or Japan + East Asia) did not accurately predict its currently invaded areas in North America, likely because the environmental conditions across the Japan + East Asia region were not sufficiently diverse and representative, and because occurrence data were relatively sparse in that region. In contrast, models that also included the European invaded range in the calibration (Japan + East Asia + Europe) successfully predicted the present distribution of *A. leucopoda* in the United States and Canada. Preliminary genetic analyses conducted by Martel et al. [8] suggested that the North American invasion likely originated from populations in Europe rather than directly from the native East Asian range. Our findings underscore the importance of incorporating both native and already-invaded regions in model calibration. Relying solely on the native range tends to underestimate the range of suitability and thus the potential areas of the establishment [66–70]. Additionally, the stage of invasion plays a critical role in determining appropriate calibration areas, as models based on early-stage invasions, with the species in question not yet at distributional equilibrium, tend to be less accurate than those calibrated under conditions closer to equilibrium [71].

Our models calibrated with the inclusion of European-invaded areas also suggested that suitable environmental conditions and host availability exist throughout the eastern United States and parts of Canada. The potential distribution of the sawfly appears to be limited to the northern and southeastern United States, extending westward only up to the Rocky Mountains and the Great Plains. These findings are consistent with studies from invaded areas in Central Europe, where elevations >580 m above sea level restrict the species’ spread [12]. The Rocky Mountains and Great Plains may serve as natural barriers to invasion, as has also been proposed for other invasive species in North America [35].

The invasion, expansion, and establishment of *A. leucopoda* across major parts of North America could have significant consequences for both native and introduced *Ulmus* populations. These iconic trees, *U. americana* in particular, have long played an important role in urban planning in the United States in view of their aesthetic appeal, resilience, and cultural symbolism [24]. Widely planted in urban settings and used as living fences in rural areas [72,73], *Ulmus* species contribute far more than visual and cultural value. As such, management efforts for *U. americana* populations amid their significant decline have a long history, aiming to mitigate ecological and potential broader impacts [74]. Although no specific estimates exist for the economic impact of *A. leucopoda*, some approaches can be used to approximate the cost of losing an urban forest (also see Hauer et al, 2020). For example, the compensation value, defined as the cost to replace or compensate an owner for a dead tree, has been estimated at 2.4 trillion dollars across 48 U.S. states [75]. Although this estimate is rough and not specific to *Ulmus* in the potentially affected states, it underscores the importance of including economic impacts on urban forests in the risk assessment of *A. leucopoda* invasions.

With the arrival of Dutch elm disease in North America in the 1930s, elm populations in both urban and natural areas experienced a drastic decline, with an estimated loss of >70 million trees [76,77]. The disease has been most prevalent in larger and older elms [78]. In this context, the invasion of *A. leucopoda*, which feeds on both young and mature elms [7], poses an additional threat that could exacerbate impacts of Dutch elm disease on these populations, especially on mature trees [7]. The negative effects of *A. leucopoda* could also have significant ecological consequences, as >500 animal species, including pollinators and herbivores, depend on *Ulmus* for habitat and resources [23].

## Conclusions

This study represents a first effort to model the potential distribution of *A. leucopoda* in North America, offering valuable insights into the areas at risk of invasion and the potential ecological and economic impacts of this species. Our models predict suitable conditions across parts of the eastern United States and Canada, while also suggesting that the Rocky Mountains and Great Plains may serve as natural barriers to its spread. The invasion of *A. leucopoda* poses an additional threat to elm populations already compromised by Dutch elm disease, with possible consequences for ecosystems, urban planning, economy, and public health. This research underscores the importance of ecological niche modeling as a critical tool for invasive species risk assessment that supports efforts in prevention, monitoring, and management. Our findings contribute to the broader understanding of invasion dynamics and provide a framework applicable to similar challenges posed by invasive species globally.

## Acknowledgments

This study has benefited greatly from discussion and commentary from our colleagues in the KUENM group, and particularly from Nikki Lemus.

## Supplementary Materials

Table S1. Summary of principal component analysis: Standard deviations, Proportion of variance explained, and Cumulative variance for each principal component.

Table S2. Area (km^2^) in North America classified as suitable for *Aproceros leucopoda* (sawfly), *Ulmus* spp., and both species.

Figure S1. Areas of potential suitability of *Aproceros leucopoda* for models with no extrapolation

Figure S2. Areas of potential suitability of *Aproceros leucopoda* for models with extrapolation and clamping

